# Severe infections emerge from the microbiome by adaptive evolution

**DOI:** 10.1101/116681

**Authors:** Bernadette C. Young, Chieh-Hsi Wu, N. Claire Gordon, Kevin Cole, James R. Price, Elian Liu, Anna E. Sheppard, Sanuki Perera, Jane Charlesworth, Tanya Golubchik, Zamin Iqbal, Rory Bowden, Ruth C. Massey, John Paul, Derrick W. Crook, Timothy E. A. Peto, A. Sarah Walker, Martin J. Llewelyn, David H. Wyllie, Daniel J. Wilson

## Abstract

Bacteria responsible for the greatest global mortality colonize the human microbiome far more frequently than they cause severe infections. Whether mutation and selection within the microbiome accompany infection is unknown. We investigated *de novo* mutation in 1163 *Staphylococcus aureus* genomes from 105 infected patients with nose-colonization. We report that 72% of infections emerged from the microbiome, with infecting and nose-colonizing bacteria showing parallel adaptive differences. We found 2.8-to-3.6-fold enrichments of protein-altering variants in genes responding to *rsp*, which regulates surface antigens and toxicity; *agr*, which regulates quorum-sensing, toxicity and abscess formation; and host-derived antimicrobial peptides. Adaptive mutations in pathogenesis-associated genes were 3.1-fold enriched in infecting but not nose-colonizing bacteria. None of these signatures were observed in healthy carriers nor at the species-level, suggesting disease-associated, short-term, within-host selection pressures. Our results show that infection, like a cancer of the microbiome, emerges through spontaneous adaptive evolution, raising new possibilities for diagnosis and treatment.

**One Sentence Summary:** Life-threatening *S. aureus* infections emerge from nose microbiome bacteria in association with repeatable adaptive evolution.

## Main Text

Communicable diseases remain a leading cause of global mortality, with bacterial pathogens among the greatest concern (1). However, many of the bacteria imposing the greatest burden of mortality, such as *Staphylococcus aureus*, are frequently found as commensal components of the body’s microbiome (2). For them invasive disease is a relatively uncommon event that is often unnecessary (3,4), and perhaps disadvantageous (5), for onward transmission. Genomics is shedding light on important bacterial traits such as host-specificity, toxicity and antimicrobial resistance (6–10). These approaches offer new opportunities to understand the role of genetics and within-host evolution in the outcome of human interactions with major bacterial pathogens (11).

Several lines of evidence support a plausible role for within-host evolution influencing the virulence of bacterial pathogens. Common bacterial infections, including *S. aureus*, are often associated with colonization of the microbiome by a genetically similar strain (12). Genome sequencing suggests that bacteria mutate much more quickly than previously accepted, and this confers a potent ability to adapt, for example evolving antimicrobial resistance *de novo* within individual patients (13,14). Opportunistic pathogens infecting cystic fibrosis patients have been found to rapidly adapt to the lung environment, with strong evidence of parallel evolution across patients (15–19). However, the selection pressures associated with antimicrobial resistance and opportunistic infections of cystic fibrosis patients may not typify within-host adaptation in common commensal pathogens that have co-evolved with humans for thousands or millions of years (20,21).

Candidate gene studies have demonstrated that certain regions, notably quorum-sensing systems such as the *S. aureus* accessory gene regulator (*agr*), mutate particularly quickly *in vivo* and in culture (22). The *agr* operon encodes a pheromone that coordinates a shift at higher cell densities from production of surface proteins promoting biofilm formation to production of secreted toxins and proteases promoting inflammation and dispersal (23). Mutants typically produce the pheromone but no longer respond to it (24). The evolution of *agr* has been variously ascribed to directional selection (25), balancing selection (26), social cheating (27) and life-history trade-off (28). However, the role of *agr* mutants in disease remains unclear, since they are frequently sampled from both asymptomatic carriage and severe infections (24).

Whole-genome sequencing case studies add weight to the idea that within-host evolution plays an important role in infection. In one persistent *S. aureus* infection, a single mutation was sufficient to permanently activate the stringent stress response, reducing growth, colony size and experimentally measured disease severity (29). In another patient, we found that bloodstream bacteria differed from those initially colonizing the nose by several mutations including loss-of-function of the *rsp* regulator (30). Functional follow-up revealed that the *rsp* mutant expressed reduced toxicity (31), but maintained the ability to cause disseminated infection (32). Unexpectedly, we found that bloodstream-infecting bacteria exhibit lower toxicity than nose-colonizing bacteria more generally (31). These results raise the question: are unique hallmarks of *de novo* mutation and selection associated with bacterial evolution in severely infected patients?

We addressed this question by investigating the genetic variants arising from within-patient evolution of *S. aureus* sampled from 105 patients with concurrent nose colonization and blood or deep tissue infection. We annotated variants to test for systematic differences between colonizing and infecting bacteria. We discovered several groups of genes showing significant enrichments of protein-altering variants indicating adaptive evolution. For genes implicated in pathogenesis, adaptive mutants were limited to infecting bacteria, while other pathways showed adaptation in the nose and infection site. Adaptive enrichments were not observed in asymptomatic carriers, nor between unrelated bacteria, indicating evolution in response to disease-associated, within-host selection pressures. Our results reveal that adaptive evolution of genes involved in regulation, toxicity, abscess formation, cell-cell communication and bacterial-host interaction drives parallel differentiation between commensal constituents of the nose microbiome and invasive infections, providing new insights into the evolution of disease in a major pathogen.

## Results

### Infecting bacteria are typically descended from the patient’s microbiome

We identified 105 patients suffering severe *S. aureus* infections admitted to hospitals in Oxford and Brighton, England, for whom we could recover contemporaneous nose swabs from admission screening. Of the 105 patients, 55 had bloodstream infections, 37 had soft tissue infections and 13 had bone and joint infections (Table 1). The infection was most often sampled on the same day as the nose, with an interquartile range of 1 day earlier to 2 days later (Table S1).

**Table 1.** Distribution of infection types and relatedness of nose-colonizing and infecting *S. aureus* among 105 patients revealed by genomic comparison.

| Infection sites | Relation of colonizing to infecting bacteria |  |  |
| --- | --- | --- | --- |
| | Unrelated<br>($\geq 1104$<br>variants) | Closely related<br>($\leq 66$ variants) | |
|  |  | Zero shared<br>genotypes | One shared<br>genotype |
| Bloodstream | 4 | 43 | 8 |
| Soft tissue | 4 | 23 | 10 |
| Bone & joint | 2 | 8 | 3 |
| Total | 10 | 74 | 21 |

To discover *de novo* mutations within and between the nose microbiome and infection site, we whole-genome sequenced 1163 bacterial colonies, a median of 5 per site. We detected single nucleotide polymorphisms (SNPs) and short insertions/deletions (indels) using previously developed, combined reference-based mapping and *de novo* assembly approaches (30,33,34). We identified 35 distinct strains, defined by multilocus sequence type (ST), across patients (Table S1). As expected (12), colonizing and infecting bacteria were usually extremely closely related (95 patients), sharing the same ST and differing by 0-66 variants. Unrelated colonizing and infecting bacteria (10 patients) differed by 1104-50573 variants and typically possessed distinct STs (e.g. Fig. 1a, Fig. S1). After excluding variants differentiating unrelated STs, we catalogued 1322 *de novo* mutations within the 105 patients.

**Fig. 1.**
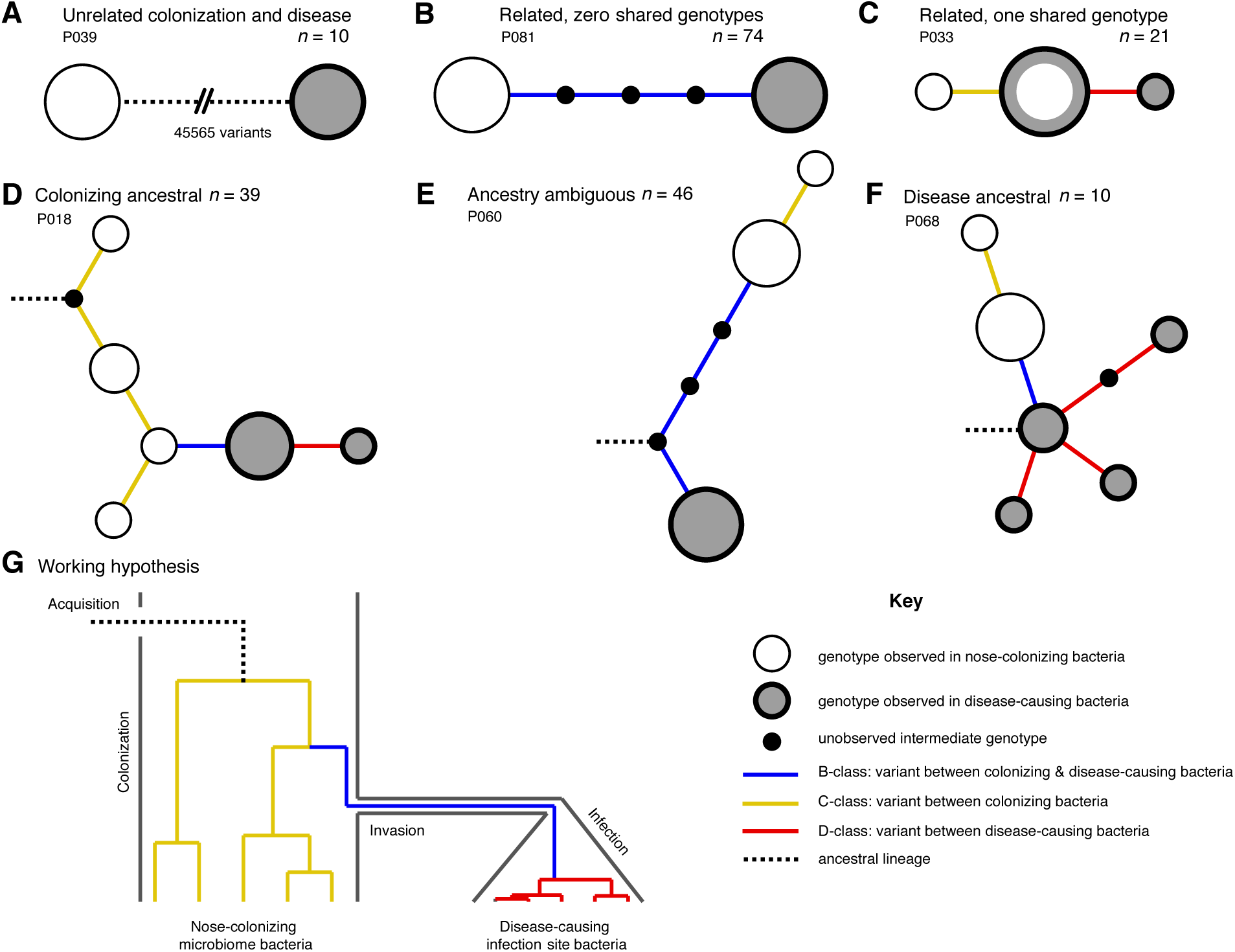
Disease-causing *S. aureus* form closely related but distinct populations descended from microbiome-colonizing bacteria in the majority of infections. Bacteria sampled from the nose and infection site of 105 patients formed one of three population structures, illustrated with example haplotrees: (**A**) Unrelated populations differentiated by many variants. (**B**) Highly related populations separated by few variants. (**C**) Highly related populations with one genotype in common. Reconstructing the ancestral genotype in each patient helped identify the ancestral population: (**D**) Nose-colonizing bacteria ancestral. (**E**) Ambiguous ancestral population. (**F**) Disease-causing bacteria ancestral. (**G**) Phylogeny illustrating the working hypothesis that variants differentiating highly related nose-colonizing and disease-causing bacteria would be enriched for variants that promote, or are promoted by, infection. In **A-F**, haplotree nodes represent observed genotypes sampled from the nose (white) or infection site (grey), with area proportional to genotype frequency, or unobserved intermediate genotypes (black). Edges represent mutations. Patient identifiers and sample sizes (*n*) are given. In **A-G**, edge color indicates that mutations occurring on those branches correspond to B-class variants between nose-colonizing and disease-causing bacteria (blue), C-class variants among nose-colonizing bacteria (gold) or D-class variants among disease-causing bacteria (red). Black dashed edges indicate ancestral lineages.

In patients with closely related strains, the within-patient population structure was always consistent with a unique migration event from the nose to the infection site, or occasionally, vice versa. Infecting and colonizing bacteria usually formed closely-related but distinct populations with no shared genotypes (74/95 patients, e.g. Fig. 1b), separated by a mean of 5.7 variants. There was never more than one identical genotype between nose-colonizing and infecting bacteria, (21/95 patients, e.g. Fig. 1c), indicating that the migration event from one population to the other involved a small number of founding bacteria (35,36). In such patients, the shared genotype likely represents the migrating genotype itself. Population structure did not differ significantly between infection types (*p* = 0.38, Table 1). Genetic diversity in the nose (mean pairwise distance, *π* = 2.8 variants) was similar to that previously observed in asymptomatic nasal carriers (33) (Reference Panel I, *π* = 4.1, *p* = 0.13), but was significantly lower in the infection site (*π* = 0.6, *p* = 10^−10.0^), revealing limited diversification post-infection.

In most patients the infection appeared to be descended from the nose. We used 1149 sequences from other patients and carriers (Reference Panel II) to reconstruct the most recent common ancestor (MRCA) for the 95/105 (90%) patients with related nose-colonizing and infecting bacteria. We thereby distinguished wild type from mutant alleles. In 49 such patients, we could determine the ancestral population. The nose microbiome was likely ancestral in 39/49 (80% of patients with related strains, or 72% of all patients) because all infecting bacteria shared *de novo* mutations in common that distinguished them from the MRCA, whereas nose-colonizing bacteria did not. In 16 of those, confidence was high because both mutant and ancestral alleles were observed in the nose, confirming it as the origin of the *de novo* mutation (e.g. Fig. 1d). Conversely, in 10/49 patients, bacteria colonizing the microbiome were likely descended from blood or deep tissue infections (20% of patients with related strains, or 18% of all patients) (e.g. Fig. 1f). Confidence was high for just three of those patients, and they showed unusually high diversity (Supplementary data, P063, P072, P093), suggesting that in persistent infections, infecting bacteria can recolonize the nose.

### Protein-truncating mutants are over-represented within infected patients

To help identify variants that could promote, or be promoted by, infection of the blood and deep tissue by bacteria colonizing the nose, we reconstructed within-patient phylogenies and classified variants by their position in the phylogeny. Sequencing multiple colonies per site enabled us to classify variants into those representing genuine differences *between* nose-colonizing and infection populations (*B*-class), variants specific to the nose-*colonizing* microbiome population (*C*-class) and variants specific to the *disease*-causing infection population (*D*-class). We hypothesized that B-class variants would be most enriched for variants promoting, or promoted by, infection, if such variants occur (Fig. 1g).

We cross-classified variants by their predicted functional effect: synonymous, non-synonymous or truncating within protein-coding sequences, or non-coding (Table 2, Table S2). As expected, the prevailing tendency of selection within patients was to conserve protein sequences, with *d*_*N*_/*d*_*S*_ ratios indicating rates of non-synonymous change 0.55, 0.68 and 0.63 times that expected under neutral evolution for B, C and D-class variants respectively.

**Table 2.** Cross-classification of variants within patients by phylogenetic position and predicted functional effect, and comparison to asymptomatic carriers. Neutrality indices (37) are defined as the odds ratio of mutation counts relative to synonymous variants in patients versus asymptomatic carriers (Reference Panel I). Those significant at *p* < 0.05 and *p* < 0.005 are emboldened and underlined respectively.

| Phylogenetic position | Number of variants (Neutrality index) |  |  |  |  |
| --- | --- | --- | --- | --- | --- |
|  | Synonymous | Non-synonymous | Protein truncating | Non-coding | Total |
| Patients with severe infections ($n=105$) | | | | | |
| Between colonization and disease (B-class) | 93 | 265 (1.1) | **39 (3.1)** | 140 (1.2) | 537 |
| Within colonization (C-class) | 93 | 325 (1.3) | **<u>59 (4.7)</u>** | 145 (1.3) | 622 |
| Within-disease (D-class) | 26 | 82 (1.2) | **15 (4.3)** | 40 (1.3) | 163 |
| Total | 212 | 672 (1.2) | **<u>113 (3.9)</u>** | 325 (1.3) | 1322 |
| Asymptomatic carriers (33) (Reference panel I, for comparison, $n=13$) | | | | | |
| Within colonization (C-class) | 37 | 97 | 5 | 45 | 184 |

In a longitudinal study of one long-term carrier, we previously reported that a burst of protein-truncating variants punctuated the transition from asymptomatic carriage to invasive infection (30). Here we found a 3.9-fold over-abundance of protein-truncating variants of all phylogenetic classes in infected patients compared to asymptomatic carriers (Reference Panel I, *p* = 0.002, Table 2), supporting the conclusion that loss-of-function mutations are disproportionately associated with evolution within infected patients. This may reflect a reduction in the efficiency with which selection removes deleterious protein-truncating mutations, and provides evidence of a systematic difference in selection within severely infected patients.

### Quorum sensing and cell-adhesion proteins exhibit adaptive evolution between colonizing and infecting bacteria

We hypothesized that variants associated with invasive infection would be enriched among the protein-altering B-class variants between the nose and infection site (Fig. 1g). Therefore we aggregated mutations by genes in a well-annotated reference genome, MRSA252, and tested each gene for an excess of non-synonymous and protein-truncating B-class variants, taking into account the length of the gene. Aggregating by gene was necessary because 1318/1322 variants were unique to single patients. The two exceptions involved non-coding variants arising in two patients each, one B-class variant 130 bases upstream of *azlC*, an azaleucine resistance protein (SAR0010), and one D-class variant 88 bases upstream of *eapH*1, a secreted serine protease inhibitor (SAR2295) (38).

We found a significant excess of five protein-altering B-class variants representing a 58.3-fold enrichment in *agrA*, which encodes the response regulator that mediates activation of the quorum sensing system at high cell densities (*p*=10^−7.5^, Fig. 2a, Table 3). The *clfB* gene encoding clumping factor B, which binds human fibrinogen and loricrin (39), showed an excess of five protein-altering B-class variants, representing a 15.9-fold enrichment that was near genome-wide significance after multiple testing correction (*p*=10^−4.7^).

**Fig. 2.**
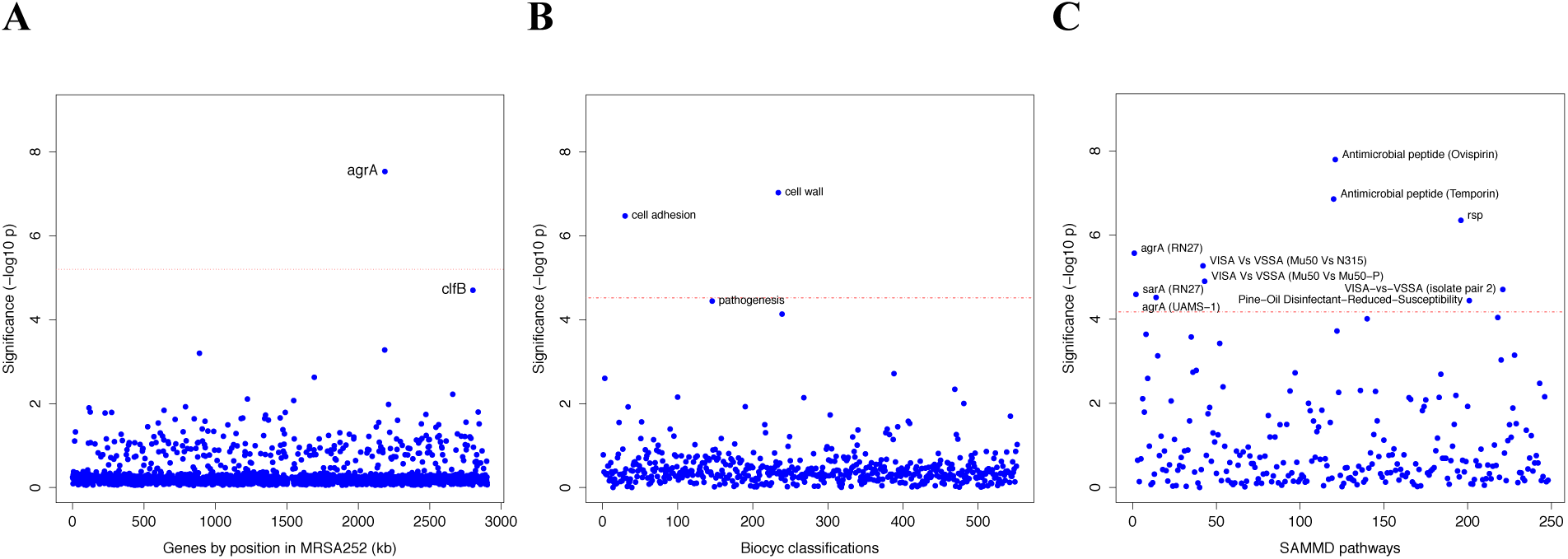
Genes, ontologies and pathways enriched for protein-altering substitutions between nose-colonizing and disease-causing bacteria within infected patients. (**A**) Significance of enrichment of 2650 individual genes. (**B**) Significance of enrichment of 552 gene sets defined by BioCyc gene ontologies. (**C**) Significance of enrichment of 248 gene sets defined by SAMMD expression pathways. Genes, pathways and ontologies that approach or exceed a Bonferroni-corrected significance threshold of *α* = 0.05, weighted for the number of tests per category, (red lines) are named.

**Table 3.** Genes, gene ontologies and expression pathways exhibiting the most significant enrichments or depletions of protein-altering B-class variants separating nose microbiome and infection site bacteria. Enrichments below one represent depletions. The total number of variants and genes available for analysis differed by database. A -log_10_ *p*-value above 5.2, 4.5 or 4.2 was considered genome-wide significant for loci, gene ontologies or expression pathways respectively (in bold).

| Gene group | No. protein-altering<br>B-class variants | | Cumulative length<br>of genes (kb) | | Enrichment | | Significance<br>($-\log_{10} p$) |
| --- | --- | --- | --- | --- | --- | --- | --- |
| Locus |  |  |  |  |  |  |  |
| *agrA* | 5 |  | 0.7 |  | 58.27 |  | **7.53** |
| *clfB* | 5 |  | 2.6 |  | 15.87 |  | 4.70 |
| Total | 289 |  | 2363.8 |  |  |  |  |
| BioCyc Gene Ontology (41) |  |  |  |  |  |  |  |
| Cell wall | 18 |  | 30.9 |  | 5.02 |  | **7.03** |
| Cell adhesion | 13 |  | 17.2 |  | 6.44 |  | **6.47** |
| Pathogenesis | 31 |  | 112.5 |  | 2.41 |  | 4.44 |
| Total | 288 |  | 2359.3 |  |  |  |  |
| SAMMD Expression Pathway | *Down-regulated* | *Up-regulated* | *Down-regulated* | *Up-regulated* | *Down-regulated* | *Up-regulated* |  |
| Ovispirin-1 (43) | 40 | 7 | 121.2 | 142.9 | 2.65 | 0.39 | **7.80** |
| Temporin L (43) | 42 | 14 | 125.1 | 156.1 | 2.78 | 0.74 | **6.86** |
| *rsp* (44) | 27 | 1 | 61.1 | 13.7 | 3.61 | 0.60 | **6.35** |
| *agrA* (RN27) (45) | 9 | 30 | 41.0 | 85.0 | 1.83 | 2.94 | **5.57** |
| VISA-vs-VSSA (Mu50 vs N315) (46) | 0 | 17 | 0 | 34.4 | 0 | 3.95 | **5.27** |
| VISA-vs-VSSA (Mu50 vs Mu50-P) (46) | 0 | 17 | 0 | 36.7 | 0 | 3.70 | **4.90** |
| VISA-vs-VSSA (isolate pair 2) (47) | 14 | 3 | 26.9 | 59.7 | 4.06 | 0.39 | **4.71** |
| *sarA* (RN27) (45) | 6 | 23 | 49.9 | 57.7 | 0.97 | 3.22 | **4.59** |
| *agrA* (UAMS-1 OD 1.0) (48) | 0 | 5 | 0 | 2.7 | 0 | 14.57 | **4.52** |
| Pine-Oil Disinfectant-Reduced-Susceptibility (49) | 17 | 5 | 36.4 | 23.6 | 3.76 | 1.70 | **4.44** |
| Total | 275 |  | 2093.5 |  |  |  |  |

Previously we identified a truncating mutation in the transcriptional regulator *rsp* to be the most likely candidate for involvement in the progression to invasive disease in one long-term nasal carrier (30). Although we observed just one variant in *rsp* among the 105 patients (3.9-fold enrichment, *p*=0.27), we found it was a non-synonymous B-class variant resulting in an alanine to proline substitution in the regulator’s helix-turn-helix DNA binding domain. In separately published experiments (32), we demonstrated that this and the original mutation induce similar loss-of-function phenotypes which, like *agr* loss-of-function mutants, express reduced toxicity, but maintained an ability to persist, disseminate and cause abscesses *in vivo*.

We found no significant enrichments of protein-altering variants among D-class variants, but we observed a significant excess of six protein-altering C-class variants in *pbp2* which encodes a penicillin binding protein involved in cell wall synthesis (19.0-fold enrichment, *p*=10^−6.0^, Fig. S2a). Pbp2 is an important target of β-lactam antibiotics (40), revealing adaption – potentially in response to antibiotic treatment – in the nose populations of some patients.

### Genes modulated by virulence regulators and antimicrobial peptides show adaptive evolution between colonizing and infecting bacteria

To improve the sensitivity to identify adaptive evolution associated with invasive infection, we developed a gene set enrichment analysis (GSEA) approach in which we tested for enrichments of protein-altering B-class variants among groups of genes. GSEA allowed us to detect signatures of adaptive evolution in groups of related genes that were not apparent when interrogating individual genes.

We grouped genes in two different ways: by gene ontology and by expression pathway. First, we obtained a gene ontology for the reference genome from BioCyc (41), which classifies genes into biological processes, cellular components and molecular functions. There were 552 unique gene ontology groupings of two or more genes. We tested for an enrichment among genes belonging to the ontology, compared to the rest of the genes.

Second, we obtained 248 unique expression pathways from the SAMMD database of transcriptional studies (42). For each expression pathway genes were classified as up-regulated, down-regulated or not differentially regulated in response to experimentally manipulated growth conditions or expression of a regulatory gene. For each expression pathway, we tested for an enrichment in genes that were up- or down-regulated compared to genes not differentially regulated.

The most significant enrichment for protein-altering B-class variants between nose and infection sites occurred in the group of genes down-regulated by the cationic antimicrobial peptide (CAMP) ovispirin-1 (*p*=10^−7.8^), with a similar enrichment in genes down-regulated by temporin L exposure (*p*=10^−6.9^, Fig. 2c). Like human CAMPs, the animal-derived ovispirin and temporin compounds inhibit epithelial infections by killing phagocytosed bacteria and mediating inflammatory responses (43). In response to inhibitory levels of ovispirin and temporin, *agr*, surface-expressed adhesins and secreted toxins are all down-regulated. Collectively, down-regulated genes showed 2.7-fold and 2.8-fold enrichments of adaptive evolution, respectively. Conversely, genes up-regulated in response to CAMPs, including the *vraSR* and *vraDE* cell-wall operons and stress response genes (43), exhibited 0.4-fold and 0.7-fold enrichments (i.e. depletions), respectively (Table 3). Thus, genes undergoing adaptive evolution are strongly inhibited by the CAMP-mediated immune response.

Genes belonging to the cell wall ontology showed the second most significant enrichment for adaptive evolution (*p*=10^−7.0^). Genes contributing to this 5.0-fold enrichment included the immunoglobulin-binding *S. aureus* Protein A (*spa*), the serine rich adhesin for platelets (*sasA*), clumping factors A and B (*clfA, clfB*), fibronectin binding protein A (*fnbA*) and bone sialic acid binding protein (*bbp*). The latter four genes contributed to another statistically significant 6.4- fold enrichment of adaptive protein evolution in the cell adhesion ontology (*p*=10^−6.5^, Fig. 3). Therefore, there is a general enrichment of surface-expressed host-binding antigens undergoing adaptive evolution.

**Fig. 3.**
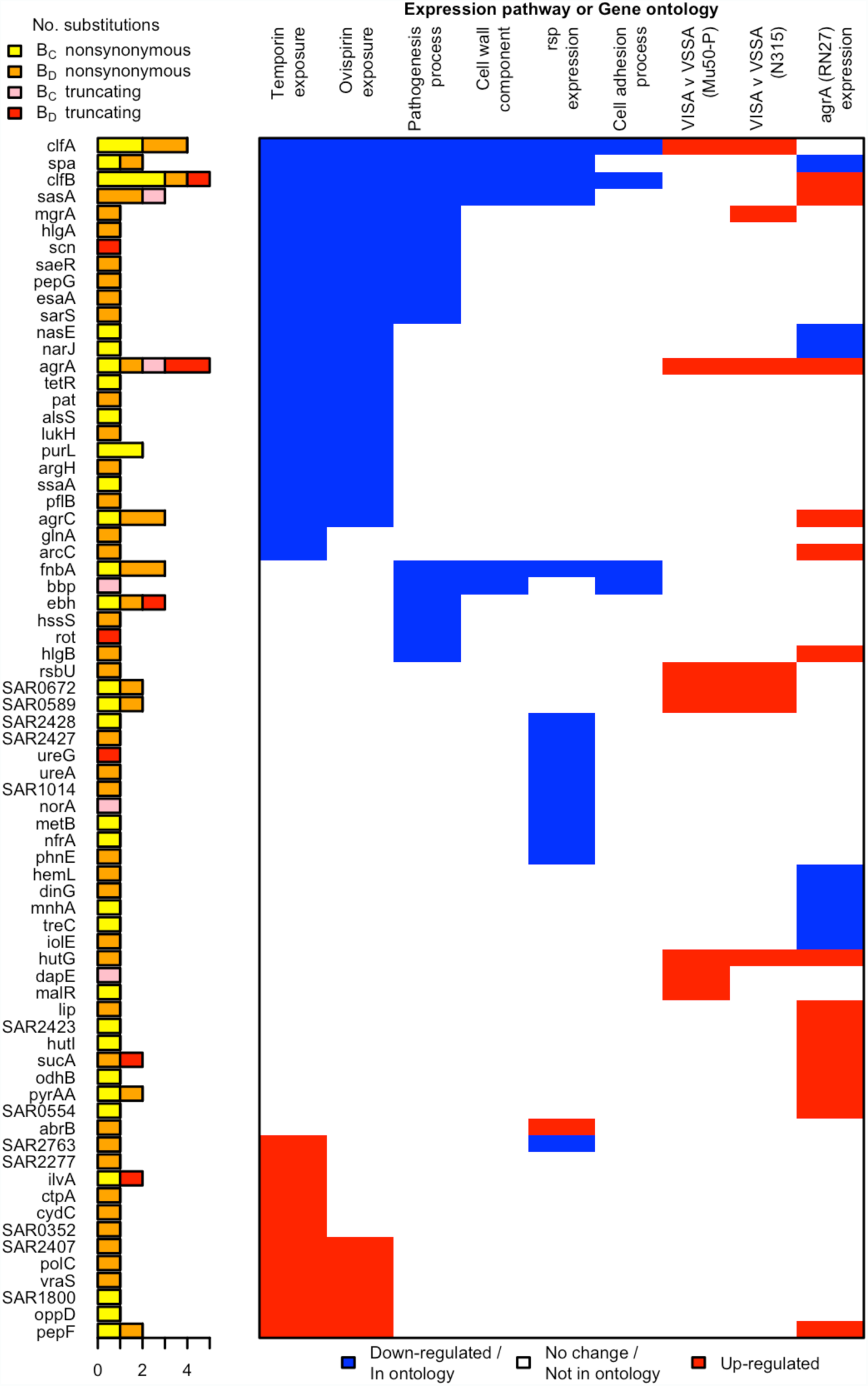
All genes contributing to the pathways and ontologies most significantly enriched for protein-altering substitutions between nose-colonizing and disease-causing bacteria. The pathogenesis ontology, in which significant enrichments were observed in disease-causing but not nose-colonizing bacteria, is shown for comparison. Every gene with at least one substitution between nose-colonizing and disease-causing bacteria and which was up-(red) or down-regulated (blue) in one of the pathways or a member of one of the ontologies (blue) is shown. To the left, the number of altering (yellow/orange) and truncating (pink/red) B-class variants is shown, broken down by the population in which the mutant allele was found: nose (B_C_; yellow/pink) or infection site (B_D_; orange/red).

The *rsp* regulon showed the most significant enrichment among gene sets defined by response to individual bacterial regulators (*p*=10^−6.4^). Genes down-regulated by *rsp* in exponential phase (44), including surface antigens and the urease operon, exhibited a 3.6-fold enrichment for adaptive evolution, while up-regulated genes showed 0.6-fold enrichment. So whereas *rsp* loss-of-function mutants were rare *per se*, genes up-regulated in such mutants were hotspots of within-patient adaptation in infected patients. Since expression is a prerequisite for adaptive protein evolution, this implies there are alternative routes by which genes down-regulated by intact *rsp* can be expressed and thereby play an important role within patients other than direct inactivation of *rsp*.

Loss-of-function in *agr* mutants represent one alternative route, since they exhibit similar phenotypes to *rsp* mutants, with reduced toxicity and increased surface antigen expression, albeit reduced ability to form abscesses (32). We found significant enrichments of genes regulated by *agrA* in two different backgrounds (*p*<10^−4.5^) and by *sarA* (*p*=10^−4.6^), underlining the influence of adaptive evolution on both secreted and surface-expressed proteins during infection. We found that expression of genes enriched for protein-altering substitutions was also altered in strains possessing reduced susceptibility to vancomycin, although not in a consistent direction across strains (*p*<10^−4.7^), and to pine-oil disinfectant (*p*=10^−4.4^), suggesting such genes may be generally involved in response to harsh environments.

Several genes contributed to multiple evolutionary signals, particularly cell-wall anchored proteins involved in adhesion, invasion and immune evasion (39), including *fnbA*, *clfA*, *clfB*, *sasA* and *spa*. These multifactorial, partially overlapping signals suggest a large target for selection in adapting to the within-patient environment (Fig. 3). The fact that we observed no comparable significant enrichments in C-class and D-class protein-altering variants (Fig. S2) indicates that these evolutionary patterns are associated specifically with the infection process.

### Adaptive evolution in pathogenesis genes is found only in infecting bacteria

Having identified adaptive evolution differentiating nose-colonizing and disease-causing bacteria, we next asked whether the mutant alleles were preferentially found in the nose or infection site. We used 1149 sequences from other patients or carriers (Reference Panel II) to reconstruct the genotype of the MRCA of colonizing and infecting bacteria respectively in each patient. This allowed us to sub-classify B-class variants by whether the mutant allele was found in the nose-colonizing bacteria (B_C_-class) or the disease-causing bacteria (B_D_-class).

*A priori*, we had expected the enrichments of adaptive evolution to be driven primarily by mutants occurring in the disease-causing bacteria (B_D_-class). One group of genes showed a signal of such an enrichment among B_D_-class variants specifically. Genes belonging to the BioCyc pathogenesis ontology were marginally genome-wide significant in B_D_-class variants, showing a 3.1-fold enrichment (*p*=10^−4.6^) and a statistically insignificant 1.7-fold enrichment in B_C_-class variants (*p*=0.13). B_D_-class mutants driving this differential signal arose in toxins including gamma hemolysin and several regulatory loci implicated in toxicity and virulence regulation: *rot*, *sarS* and *saeR*.

Surprisingly however, we found that all other significantly enriched gene sets were driven by mutant alleles occurring both in colonizing and infecting bacteria (Fig. S3). This indicates there are common selection pressures in the nose and infection site during the process of infection within patients, leading to convergent evolution across body sites. So while adaptation in pathogenesis genes appears specifically invasion-associated, other signals of adaptation in severely infected patients are driven by selection pressures, which might compensate for an altered within-host environment during infection, that are as likely to favor mutants in nose-colonizing bacteria as infecting bacteria.

### Signals of adaptation are specific to infected patients and differ from prevailing signatures of selection

Two lines of evidence show that the newly discovered signatures of within-host adaptive evolution, both in infecting and nose-colonizing bacteria, are unique to evolution in infected patients. To test this theory against the alternative explanation that our approach merely detects the most rapidly evolving proteins, we searched for similar signals in alternative settings: evolution within asymptomatic carriers, and species-level evolution between unrelated bacteria.

There was no significant enrichment of protein-altering variants in any gene, ontology or pathway among 235 variants identified from 10 longitudinally sampled asymptomatic nasal carriers (Reference Panel III, Fig. S4, Table S3). To address the modest sample size, we performed goodness-of-fit tests, focusing on the signals most significantly enriched in patients. We found significant depletions of protein-altering variants in carriers relative to patients in the *rsp*, *agr* and *sarA* regulons (*p*=10^−4.0^) and the pathogenesis ontology (*p*=10^−3.2^, Table S4).

Nor were the relative rates of non-synonymous to synonymous substitution (*d*_*N*_/*d*_*S*_) higher between unrelated *S. aureus* (Reference Panel IV) in the genes that contributed most to the signals associated with adaptation within patients: *agrA*, *agrC clfA*, *clfB*, *fnbA* and *sasA*. Although synonymous diversity was somewhat higher than typical in these genes, the *d*_*N*_/*d*_*S*_ ratios showed no evidence for excess protein-altering change in these compared to other genes (Fig. S5). Accordingly, incorporating this locus-specific variability of *d*_*N*_/*d*_*S*_ into the GSEA did not affect the results (Fig. S6). Taken together these lines of evidence show that the ontologies, pathways and genes significantly differentiated between colonizing and infecting bacteria arise in response to selection pressures specifically associated with infected patients, and are not repeated in asymptomatic carriers or species-level evolution.

## Discussion

We have discovered that common, life-threatening infections of *S. aureus* are frequently descended from bacteria colonizing the human microbiome. These infections are associated with repeatable patterns of bacterial evolution driven by within-patient mutation and selection. Genes involved in pathogenesis, notably toxins and regulators, showed evidence for adaptation in infecting but not nose-colonizing bacteria. Surprisingly, other signatures of adaptation occurred in parallel in nose-colonizing and infecting bacteria, affecting genes responding to cationic antimicrobial peptides and the virulence regulators *rsp* and *agr*. Such genes mediate toxicity, abscess formation, immune evasion and bacterial-host binding. Adaptation within both regulator and effector genes reveals that multiple, alternative evolutionary paths are targeted by selection in infected patients.

The signatures of within-patient adaptation that we found differed from prevailing signals of selection at the species level. This discordance means that infection-associated adaptive mutations within patients are rarely transmitted, and argues against a straightforward host-pathogen arms race as the predominant evolutionary force acting within and between patients. Instead, it supports the notion of a life-history trade-off between adaptations favoring colonization and infection distinct from those favoring dissemination and onward transmission. As such, invasive disease may be analogous to cancer in multicellular organisms, representing an ever-present risk of mutations in the microbiome favored by short-term selection but ultimately incidental or damaging to the bacterial reproductive life cycle.

Nor did we see these signatures of bacterial adaptation and excess loss-of-function mutations in healthy nose carriers, indicating that risk factors for invasive infections, such as a weakened or over-activated immunological response, comorbidities or medical interventions, may create distinctive selection pressures in infected patients. As in cancer, the effects of such risk factors may be mediated, at least in part, through the selection pressure they exert on the microbiome.

The existence of signatures of adaptive substitutions associated with invasive disease raises the possibility of developing new diagnostic techniques and personalizing treatment to the individual patient’s microbiome. The ability of genomics to characterize the selective forces driving adaption within the human body in unprecedented detail provides new opportunities to improve experimental models of disease. Ultimately, it may be possible to develop therapies that utilize our new understanding of within-patient evolution to target the root causes of invasive disease from the bacterial perspective.

## Acknowledgments

We would like to thank Ed Feil, Stephen Leslie, Gil McVean and Richard Moxon for helpful insights and useful discussions. Sequencing reads uploaded to short read archive (SRA) under BioProject PRJNA369475. RNA-Seq data relating to isolate from P005 (aka ‘patient S’) previously submitted under BioProject PRJNA279958.

The views expressed in this publication are those of the authors and not necessarily those of the funders. This study was supported by the Oxford NIHR Biomedical Research Centre, a Mérieux Research Grant, the National Institute for Health Research Health Protection Research Unit (NIHR HPRU) in Healthcare Associated Infections and Antimicrobial Resistance at Oxford University in partnership with Public Health England (PHE) (grant HPRU-2012-10041), and the Health Innovation Challenge Fund (a parallel funding partnership between the Wellcome Trust (grant WT098615/Z/12/Z) and the Department of Health (grant HICF-T5-358)). T.E.P. and D.W.C. are NIHR Senior Investigators. D.J.W. and Z.I. are Sir Henry Dale Fellows, jointly funded by the Wellcome Trust and the Royal Society (Grants 101237/Z/13/Z and 102541/Z/13/Z). B.C.Y is a Research Training Fellow funded by the Wellcome Trust (Grant 101611/Z/13/Z). We acknowledge the support of Wellcome Trust Centre for Human Genetics core funding (Grant 090532/Z/09/Z).

## Supplementary Materials

### Materials and Methods

#### Patient sample collection

105 patients with severe *S. aureus* infections for whom the organism could be cultured from both admission screening nasal swab and clinical sample were identified from the microbiological laboratories of hospitals in Oxford and Brighton, England. This study design builds in robustness to potential confounders by matching disease-causing and nose-colonizing bacteria within the same patients. Clinical samples comprised 55 blood cultures and 50 pus, soft tissue, bone or joint samples. The bacteria sampled and sequenced from one patient (‘patient S’, P005 in this study) have been previously described (32). Five individuals had both blood and another culture-positive clinical sample; we focus analysis on the blood sample. Nasal swabs were incubated in 5% NaCl broth overnight at 37C, then streaked onto SASelect agar (BioRad) and incubated overnight at 37C. We picked five colonies per sample (twelve during the pilot phase involving nine patients), streaked each onto Columbia blood agar and incubated overnight at 37C for DNA extraction. Clinical samples were handled according to the local laboratory standard operating procedure for pus, sterile site and blood cultures. When bacterial growth was confirmed as *S. aureus*, the primary culture plate was retrieved, and multiple colonies were picked. These were streaked onto Columbia blood agar and incubated overnight at 37C for DNA extraction. Sequencing multiple colonies per site allowed us to distinguish genuine genetic differences between nose-colonizing and disease-causing bacteria from transient variants.

#### Reference Panels

For comparison to the patient-derived isolates, we collated previously described samples from other sources to construct four Reference Panels: I. A collection of 131 genomes capturing cross-sectional diversity in the noses of 13 asymptomatic carriers (33), arising from a carriage study of *S. aureus* in Oxfordshire (50) (BioProject PRJEB2881). II. A compilation of 95 unrelated samples from the same Oxfordshire carriage study (BioProject accession number PRJEB255), 145 sequences from a study of within-host evolution of *S. aureus* in 3 individuals (30) (BioProject PRJEB2892) and 909 sequences from nasal carriage and bloodstream infection used in a study of whole genome sequencing to predict antimicrobial resistance (51) (BioProject PRJEB6251). We used these samples to improve our reconstruction of ancestral genotypes in each patient. III. A collection of 237 genomes from longitudinal samples from 10 patients (33,52), (BioProject PRJEB2862) arising from the same Oxfordshire carriage study. We used these to compare evolution within patients and asymptomatic carriers. IV. A collection of 16 previously-published high-quality closed reference genomes, comprising unrelated isolates mainly of clinical and animal origin: MRSA252 (Genbank accession number BX571856.1), MSSA476 (BX571857.1), COL (CP000046.1), NCTC 8325 (CP000253.1), Mu50 (BA000017.4), N315 (BA000018.3), USA300_FPR3757 (CP000255.1), JH1 (CP000736.1), Newman (AP009351.1), TW20 (FN433596.1), S0385 (AM990992.1), JKD6159 (CP002114.2), RF122 (AJ938182.1), ED133 (CP001996.1), ED98 (CP001781.1), EMRSA15 (HE681097.1) (53–65). We used these to contrast species-level evolution to within-patient evolution.

#### Whole genome sequencing

For each bacterial colony, DNA was extracted from the subcultured plate using a mechanical lysis step (FastPrep; MPBiomedicals, Santa Ana, CA) followed by a commercial kit (QuickGene; Fujifilm, Tokyo, Japan), and sequenced at the Wellcome Trust Centre for Human Genetics, Oxford on the Illumina (San Diego, California, USA) HiSeq 2000 platform, with paired-end reads 101 base pairs for 9 patients in the pilot phase, and 150 bases in the remainder.

#### Variant calling

We used Velvet (66) to assemble reads into contigs *de novo*, and Stampy (67) to map reads against two reference genomes: MRSA252 (53) and a patient-specific reference comprising the contigs assembled for one colony sampled from each patient’s nose. Repetitive regions, defined by BLASTing (68) the reference genome against itself, were masked prior to variant calling. To obtain multilocus sequence types (69) we used BLAST to find the relevant loci, and looked up the nucleotide sequences in the online database at http://saureus.mlst.net/.

Bases called at each position in the reference and those passing previously described (30,33,70) quality filters were used to identify single nucleotide polymorphisms (SNPs) from Stampy-based mapping to MRSA252 and the patient-specific reference genomes. We used Cortex (34) to identify SNPs and short indels. Variants found by Cortex were excluded if they had fewer than ten supporting reads or if the base call was heterozygous at more than 5% of reads.

Where physically clustered variants with the same pattern of presence/absence across genomes were found, these were considered likely to represent a single evolutionary event: tandem repeat mutation or recombination. These were de-duplicated to a single variant to avoid inflating evidence of evolutionary events in these regions.

#### Variant annotation and phylogenetic classification

Maximum likelihood trees were built to infer bacterial relationships within patients (71). To prioritize variants for further analysis, they were classified according to their phylogenetic position in the tree: B-class (between colonization and disease), C-class (within colonizing population) and D-class (within disease population). Variants were cross-classified by their predicted functional effect based on mapping to the reference genome or BLASTing to a reference allele: synonymous, non-synonymous or truncating for protein-coding sequences, or non-coding.

Where variation was found using a patient-specific reference, these variants were annotated by first aligning to MRSA252 using Mauve (72). If no aligned position in MRSA252 could be found, additional annotated references were used. Where variation was found using Cortex only, the variant was annotated by first locating it by comparing the flanking sequence to MRSA252 and other annotated references using BLAST. MRSA252 orthologs were identified using geneDB (73) and KEGG (74).

#### Reconstructing ancestral genotypes per patient

We constructed a species-level phylogeny for all bacteria sampled from the 105 patients together with Reference Panel II (unrelated asymptomatic carriage isolates and bacteremia isolates) using a two-step neighbor-joining and maximum likelihood approach, based on a whole-genome alignment derived from mapping all genomes to MRSA252. We first clustered individuals into seven groups using neighbour-joining (75), before resolving the relationships within each cluster by building a maximum likelihood tree using RAxML (76), assuming a general time reversible (GTR) model. To overcome a limitation in the presence of divergent sequences whereby RAxML fixes a minimum branch length that may be longer than a single substitution event, we fine-tuned the estimates of branch lengths using ClonalFrameML (77). We used these subtrees to identify, for each patient, the most closely related ‘nearest neighbor’ sampled from another patient or carrier. We employed this nearest neighbor as an outgroup, and used the tree to reconstruct the sequence of the MRCA of colonizing and infecting bacteria for each patient using a maximum likelihood method (78) in ClonalFrameML (77). This in turn allowed us to identify the ancestral (wild type) and derived (mutant) allele for all variants mapping to MRSA252. For variants not mapping to MRSA252, we repeated the Cortex variant calling analysis, including the nearest neighbor, and identified the ancestral allele as the one possessed by the nearest neighbor. This approach allowed us to identify ancestral versus derived alleles for 97% of within-patient variants. We used the reconstructions of the within-patient MRCA sequences and identity of ancestral vs derived alleles to sub-categorize B-class variants into those in which the mutant allele was found in the colonizing population (B_C_-class) versus the disease-causing population (B_D_-class). 521 (97%) of B-class variants were typeable, and in 281 (54%) of these, the mutant allele was found in the disease population. This allowed us to test for differential enrichments in these two sub-classes.

#### Mean pairwise genetic diversity

Separately for the nose site and infection site of each patient, we calculated the mean pairwise diversity *π* as the mean number of variants differing between each pair of genomes. We compared the distributions of *π* between patients and Reference Panel II (13 cross-sectionally sampled asymptomatic carriers) using a Mann-Whitney-Wilcoxon test.

#### Calculating d_N_/d_S_ ratio

For assessing the *d*_*N*_*/d*_*S*_ ratio within patients, we adjusted the ratio of raw counts of total numbers of non-synonymous and synonymous SNPs by the ratio expected under strict neutrality. We estimated that the rate of non-synonymous mutation was 4.9 times higher than that of synonymous mutation in *S. aureus* based on codon usage in MRSA252 and the observed transition:transversion ratio in non-coding SNPs.

#### The Neutrality Index

To compare the relative *d*_*N*_*/d*_*S*_ ratios between two groups of variants we computed a Neutrality Index as *R*_1_/*R*_2_ where *R*_1_ and *R*_2_ were the ratio of counts of non-synonymous to synonymous variants in each group respectively (37). We compared B, C and D-class variants within patients to C-class patients within Reference Panel I (13 cross-sectionally sampled asymptomatic carriers). A Neutrality Index in excess of one indicates a higher *d*_*N*_*/d*_*S*_ ratio in the former group. We used Fisher’s exact test to evaluate the significance of the differences between the groups.

#### Gene enrichment analysis

To test for significant enrichment of variants in a particular gene, we employed a Poisson regression in which we modelled the expected numbers of *de novo* variants across patients in any gene *j* as *λ*_0_*L*_*j*_ under the null hypothesis of no enrichment, where *λ*_0_ gives the expected number of variants per kilobase and *L*_*j*_ is the length of gene *j* in kilobases. We compared this to the alternative hypothesis in which the expected number of variants was *λ*_*i*_*L*_*i*_ for gene *i*, the gene of interest, and *λ*_1_*L*_*j*_ for any other gene *j*. Using R (79), we estimated the parameters *λ*_0_, *λ*_1_ and *λ*_*i*_ from the data by maximum likelihood and tested for significance via a likelihood ratio test with one degree of freedom. This procedure assumes no recombination within patients, which was reasonable since we found little evidence of recombination in this study or previously (33), including no within-host genetic incompatibilities, and we removed physically clustered variants associated with possible recombination events. We analysed all protein-coding genes in MRSA252, testing for an enrichment of variants expected to alter the transcribed protein (both non-synonymous and truncating mutations). These tests were also applied to synonymous mutations and no enrichments were found.

#### Gene set enrichment analysis

Since the number of genes outweighed the number of variants detected, we had limited power to detect weak to modest enrichments at the individual gene level. Instead we pooled genes using ontologies from the BioCyc MRSA252 database (41) and expression pathways from the SAMMD database of transcriptional studies (42). The BioCyc database comprises ontologies describing biological processes, cellular components and molecular functions. The SAMMD database groups genes up-regulated, down-regulated or not differentially regulated in response to experimentally manipulated growth conditions or isogenic mutations, usually of a regulatory gene. After excluding ontologies or pathways with two groups, one involving a single gene, and combining ontologies or pathways with identical groupings of genes, we conducted 800 GSEAs in addition to the 2650 ontologies comprised of individual loci. The number of groupings of genes was always two for BioCyc (included/excluded from the ontology) and two or three for SAMMD (up-/down-/un-differentially regulated in the experiment). Again we employed a Poisson regression in which we modelled the expected numbers of variants in any gene *j* as *λ*_0_*L*_*j*_ under the null hypothesis of no enrichment where *λ*_0_ gives the expected number of variants per kilobase and *L*_*j*_ is the length of gene *j* in kilobases. We compared this to the alternative hypothesis in which the expected number of variants was *λ*_1_*L*_*j*_, *λ*_2_*L*_*j*_ or *λ*_3_*L*_*j*_ for gene *j* depending on the grouping in the ontology/pathway. Using R, we estimated the parameters *λ*_0_, *λ*_1_, *λ*_2_ and *λ*_3_ from the data by maximum likelihood and tested for significance via a likelihood ratio test with one or two degrees of freedom, depending on the number of groupings in the ontology/pathway.

#### GSEA multiple testing correction

To account for the multiplicity of testing, we adjusted the *p-value* significance thresholds from a nominal *α* = 0.05 using the weighted Bonferroni method. We weighted the significance thresholds by the relative number of tests in each category: 2650 genes, 552 BioCyc ontologies and 248 SAMMD expression pathways. This avoids overly stringent multiple testing correction in categories with fewer tests (80), e.g. the 248 SAMMD expression pathways, owing to other categories with very large numbers of tests, e.g. the 2650 genes. This gave adjusted significance thresholds of 10^−5.2^ for genes, 10^−4.5^ for BioCyc ontologies and 10^−4.2^ for SAMMD expression pathways.

#### Longitudinal evolution in asymptomatic carriers

To test whether the patterns of evolution we observed between colonizing and invading bacteria in severely infected patients were typical or unusual, we analysed Reference Panel III (a collection of 10 longitudinally sampled asymptomatic carriers). Since natural selection is more efficacious over longer periods of time, the longitudinal sampling of these individuals gave us greater opportunity to detect subtle evolutionary patterns in asymptomatic carriers. We characterized variation in these carriers as in the patients. Given the modest sample size and smaller number of variants detected in these individuals (235), we performed GSEA to test for enrichments only in particular genes, ontologies and pathways that were significantly enriched within patients, requiring less stringent multiple testing correction.

#### omegaMap analysis

We estimated *d*_*N*_*/d*_*S*_ ratios between unrelated *S. aureus* to characterize the prevailing patterns of selection at the species level. We used Mauve (72) to pairwise align 15 reference genomes against MRSA252, i.e. Reference Panel IV. This allowed us to distinguish orthologs from paralogs in the next step in which we multiply aligned all coding sequences overlapping those in MRSA252 using PAGAN (81). After removing sequences with premature stop codons, we analysed each alignment of between two and 16 genes using a modification of omegaMap (82), assuming all sites were unlinked. We previously showed this assumption, which confers substantial computational efficiency savings, does not adversely affect estimates of selection coefficients (83). We estimated variation in *d*_*N*_*/d*_*S*_ within genes using Monte Carlo Markov chain, running each chain for 10,000 iterations. We assumed exponential prior distributions on the population scaled mutation rate (*θ*), the transition:transversion ratio (*κ*) and the *d*_*N*_*/d*_*S*_ ratio (*ω*) with means 0.05, 3 and 0.2 respectively. We assumed equal codon frequencies and a mean of 30 contiguous codons sharing the same *d*_*N*_*/d*_*S*_ ratio. For each gene, we computed the posterior mean *d*_*N*_*/d*_*S*_ ratio across sites. This allowed us to rank the relative strength of selection across genes in MRSA252, and to account for differences in *d*_*N*_*/d*_*S*_, as well as gene length, in the GSEA. We achieved this by modifying the expected number of variants in gene *j* to be *λ*_0_*ω*_*j*_*L*_*j*_ under the null hypothesis of no enrichment versus *λ*_1_*ω*_*j*_*L*_*j*_, *λ*_2_*ω*_*j*_*L*_*j*_ or *λ*_3_*ω*_*j*_*L*_*j*_ under the alternative hypothesis depending on the ontology or pathway, where *ω*_*j*_ is the posterior mean *d*_*N*_*/d*_*S*_ in gene *j*.

#### Ethical framework

Ethical approval for linking genetic sequences of *S. aureus* isolates to patient data without individual patient consent in Oxford and Brighton in the U.K. was obtained from Berkshire Ethics Committee (10/H0505/83) and the U.K. Health Research Agency [8-05(e)/2010].

#### Accession numbers

Sequencing reads uploaded to short read archive (SRA) under BioProject PRJNA369475. RNA-Seq data relating to isolate from P005 (aka ‘patient S’) previously submitted under BioProject PRJNA279958.

## Author contributions

BCY, study design, sample collection, DNA extraction, bioinformatics, analysis, writing. C-HW, bioinformatics, analysis, writing. NCG, JRP, sample collection, DNA extraction. KC, EL, SP, DNA extraction. AS, JC, TG, ZI, bioinformatics. RB, RCM, study design, interpretation. JP, DWC, TEAP, ASW, MJL, study design, sample collection, interpretation. DHW, study design, analysis. DJW, study design, analysis, writing.

---

**Fig. S1.**
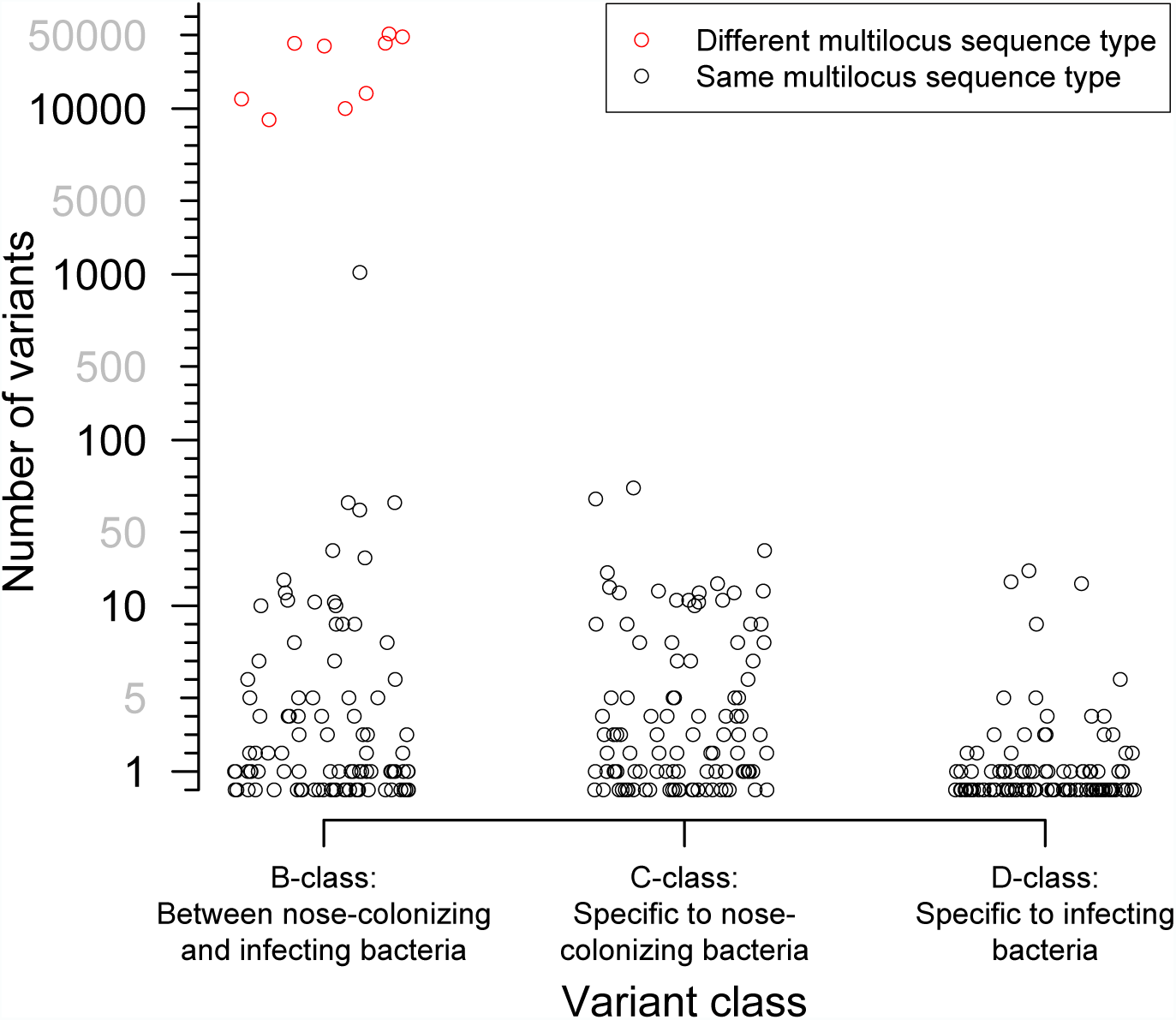
Distribution of the number of variants identified within 105 severely infected patients, by class. Three classes of variants were identified: those representing genuine differences *between* nose-colonizing and infection populations (*B*-class), variants specific to the nose-*colonizing* microbiome population (*C*-class) and variants specific to the *disease*-causing infection population (*D*-class). The number of variants is shown on a piecewise-linear axis, with horizontal positioning permuted to assist visualization. Where nose-colonizing and infecting bacteria possessed different multilocus sequence types, the number of variants between those populations is colored red. When the number of B-class variants was 66 or less, nose-colonizing and infecting bacteria were considered related, since a similar range of (C-class) diversity was observed within the nose-colonizing populations of bacteria with the same multilocus sequence type. When the number of B-class variants was 1104 or more, nose-colonizing and infecting bacteria were considered unrelated.

**Fig. S2.**
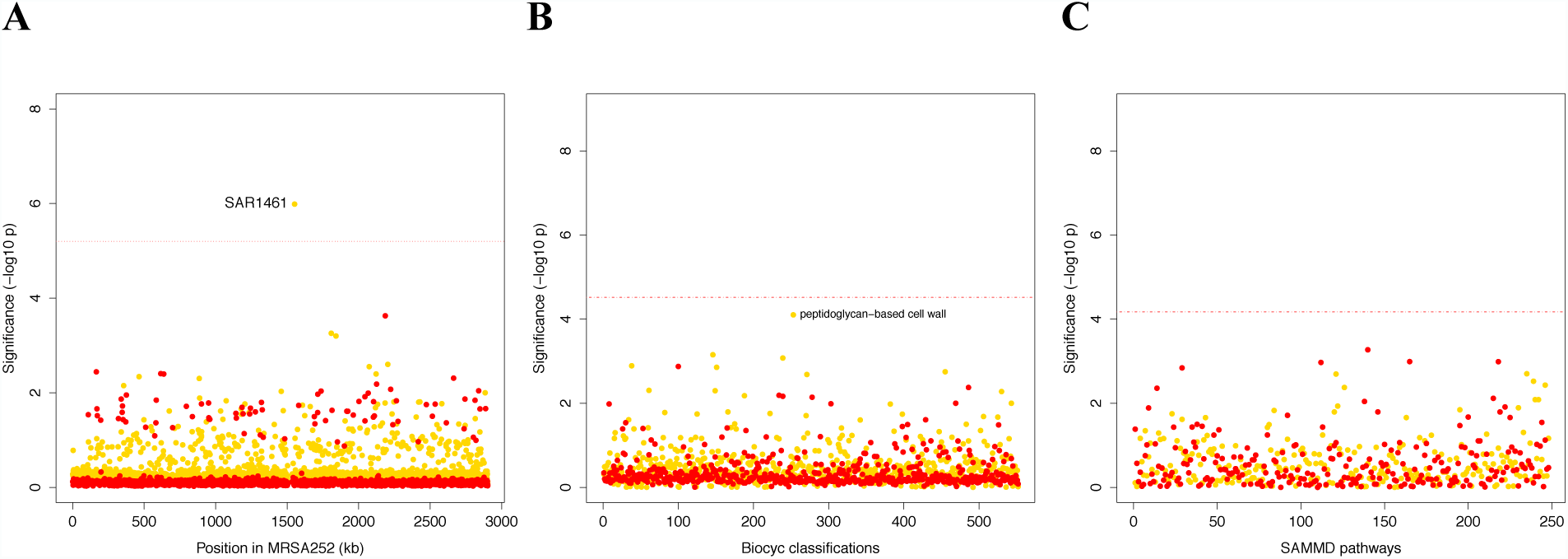
Genes, ontologies and pathways enriched for protein-altering transient variants within nose-colonizing and disease-causing bacteria. (**A**) Significance of enrichment of 2650 individual genes. SAR1461 encodes Pbp2, penicillin-binding protein 2. (**B**) Significance of enrichment of 552 gene sets defined by BioCyc gene ontologies. (**C**) Significance of enrichment of 248 gene sets defined by SAMMD expression pathways. C-class variants among nose-colonizing bacteria are colored gold, D-class variants among disease-causing bacteria are colored red. Genes, pathways and ontologies that approach or exceed a Bonferroni-corrected significance threshold of *α* = 0.05, weighted for the number of tests per category, (red lines) are named.

**Fig. S3.**
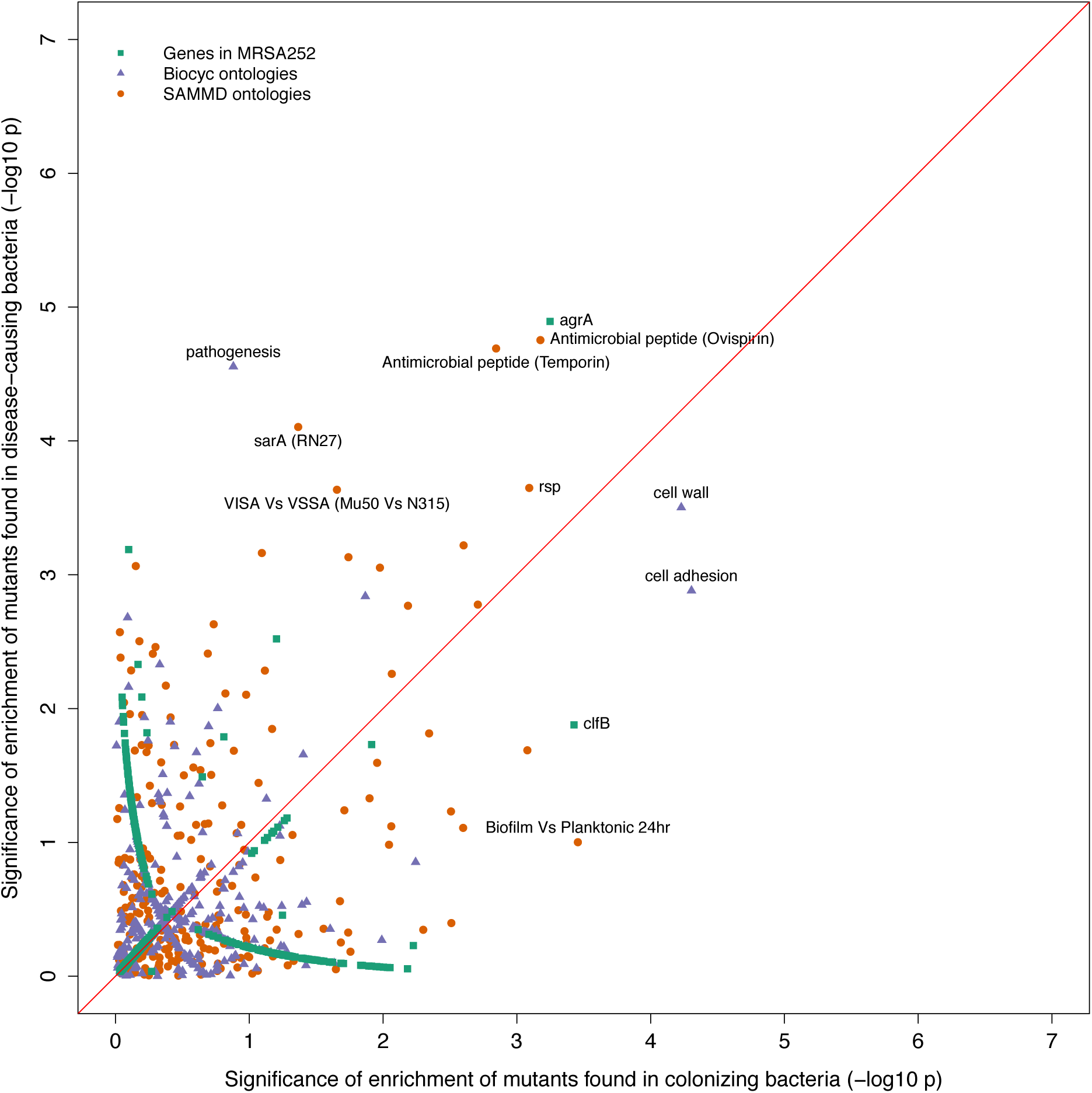
Gene set enrichment analysis of B-class mutants occurring in the nose or the infection site. Each point indicates the –log10 *p*-values of two tests for enrichment of protein-altering variants found among mutants in nose-colonizing bacteria vs disease-causing bacteria. The shape of each point represents the type of enrichment tested (squares: within 2650 genes in MRSA252, triangles: 552 BioCyc gene ontologies, circles: 248 SAMMD expression pathways). A line of 1:1 correspondence is plotted in red. A -log_10_ *p*-value above 5.2, 4.5 or 4.2 was considered genome-wide significant for loci, gene ontologies or expression pathways respectively.

**Fig. S4.**
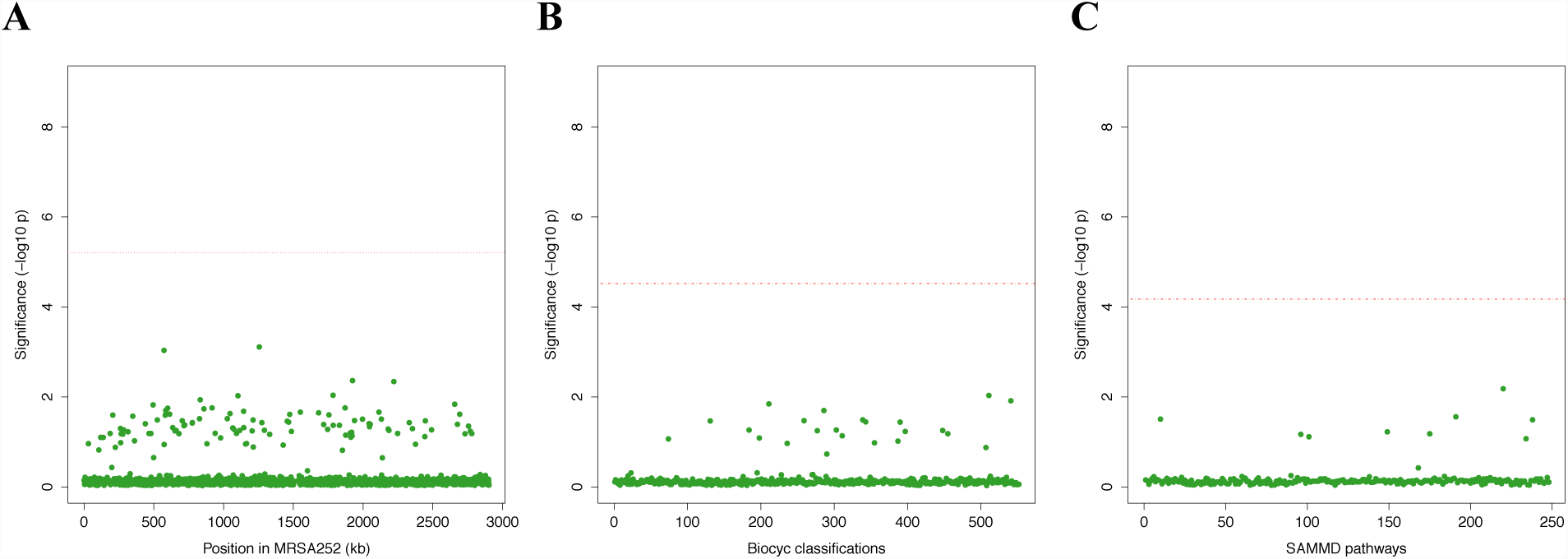
Genes, ontologies and pathways enriched for protein-altering variants among longitudinally sampled asymptomatic nasal carriers. (**A**) Significance of enrichment of 2650 individual genes. (**B**) Significance of enrichment of 552 gene sets defined by BioCyc gene ontologies. (**C**) Significance of enrichment of 248 gene sets defined by SAMMD expression pathways. Genes, pathways and ontologies that approach or exceed a Bonferroni-corrected significance threshold of *α* = 0.05, weighted for the number of tests per category, (red lines) are named.

**Fig. S5.**
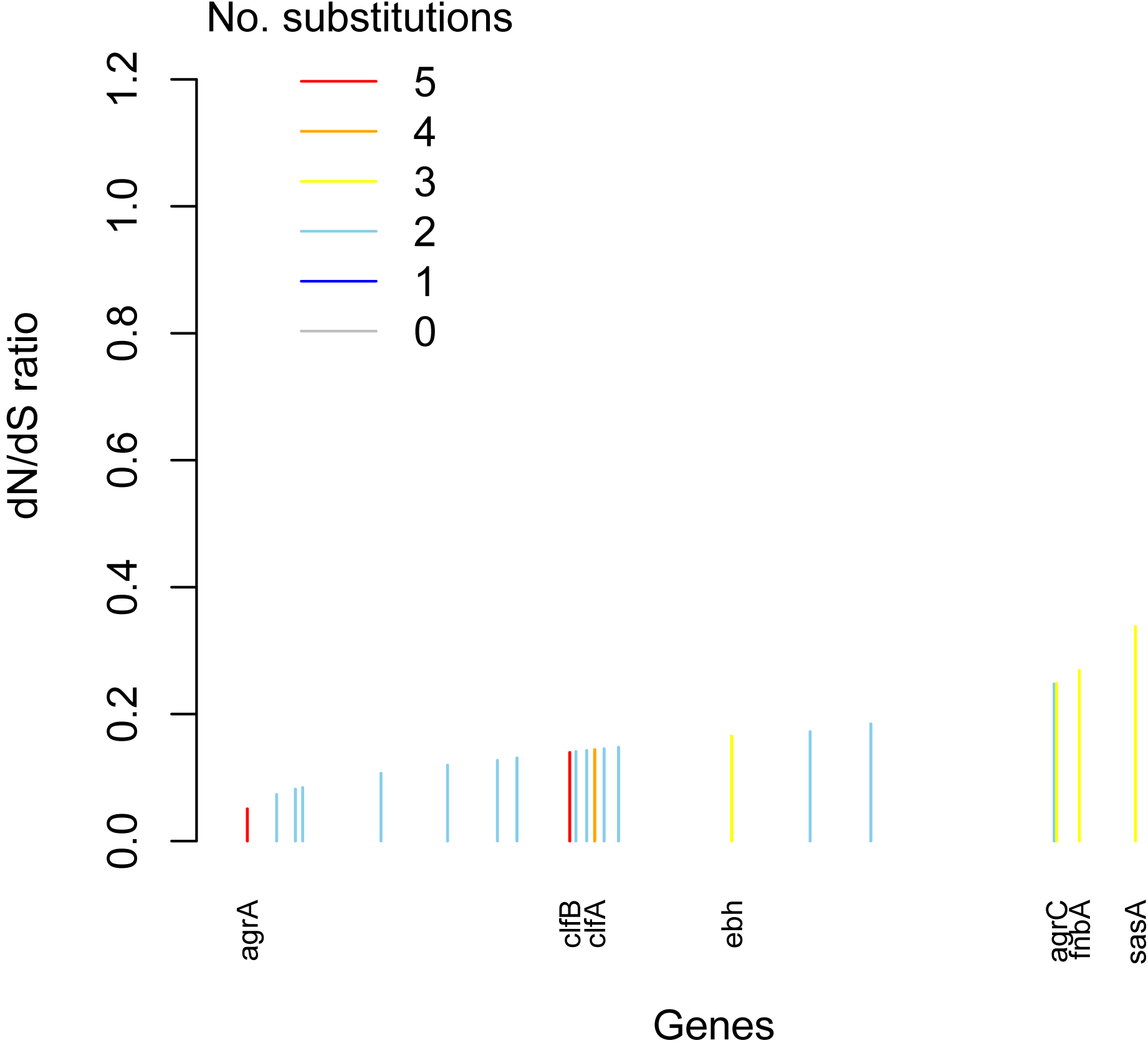
Genes enriched for substitutions between nose-colonizing and disease-causing bacteria within patients are not the most rapidly evolving at the species level. An estimate of the *d*_*N*_/*d*_*S*_ ratio between unrelated bacteria is shown for each gene, color-coded by the number of protein-altering substitutions between nose-colonizing and disease-causing bacteria within patients. There was a negative Spearman rank correlation between *d*_*N*_/*d*_*S*_ ratio and substitutions within patients (*ρ* = –0.04, *p* = 0.02).

**Fig. S6.**
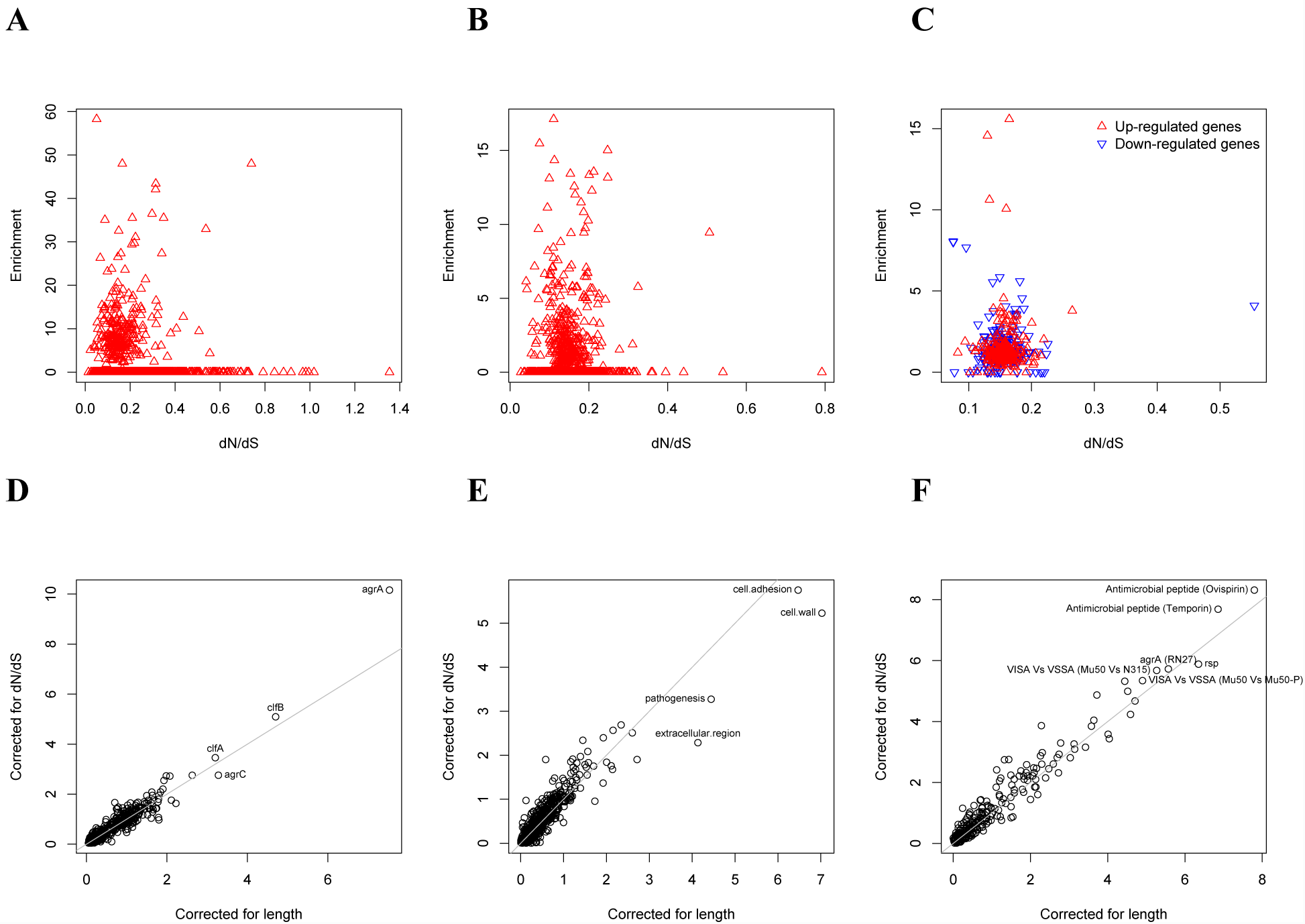
Gene set enrichment analysis is robust to species-level differences in *d*_*N*_/*d*_*S*_ between genes. For every locus, expression pathway and gene ontology, we estimated *d*_*N*_/*d*_*S*_ between unrelated *S. aureus*. There was no relationship between *d*_*N*_/*d*_*S*_ and enrichment of protein-altering substitutions between nose-colonizing and disease-causing bacteria in (**A**) loci, (**B**) ontologies nor (**C**) pathways (non-significant correlations, *p* > 0.05). When we incorporated variability in *d*_*N*_/*d*_*S*_ between genes in the gene set enrichment analyses, the results were robust for (**D**) loci, (**E**) ontologies and (**F**) pathways, showing only small differences in significance (-log_10_ *p*-value) between the analyses that correct for locus length only (horizontal axes) and those that correct for locus length and *d*_*N*_/*d*_*S*_ (vertical axes).

**Table S1**. List of all cultures included in the site, the site of infection (and any known source if bloodstream), number of isolates sequenced from each site, ST or CC by in silico MLST, number of variants found at each site and the mean pair-wise difference comparing isolates.

**Table S2**. List of all variants found within patients with *S. aureus* disease, location on shared reference (MRSA252), or position and reference genome name and accession number if variant could not be localized on MRSA252. Each variant is described by the alleles found, its location in gene, the predicted effect on gene product and the location of the variant on the phylogenetic tree.

**Table S3**. List of all variants found within long term asymptomatic carriers, location on shared reference (MRSA252), or position and reference genome name and accession number if variant was not localized on MRSA252. Each variant is described by the alleles found, its location in gene and the predicted effect on gene product.

**Table S4.** For all ontologies showing enrichment in within-patient B_D_-class variants, we identified the genes with variants contributing to the signal. We counted the number of protein-altering variants in these genes within patients, and compared to the number in long-term asymptomatic carriers. P values calculated using Fisher’s exact test. *Variant totals are different for SAMMD pathways (*rsp, agrA, sarA*) and BioCyc ontologies (cell wall, cell adhesion, pathogenesis) because pathway information is available for a different number of loci in each database.

| Gene Ontology or Expression Pathway<br>(Loci with protein-altering B<sub>D</sub>-class variants within patients) | Number of variants* |  | *p*-value |
| --- | --- | --- | --- |
|  | Within patients | Within carriers |  |
| AgrA locus (SAR2126) | 3/156 | 0/115 | n.s. |
| Rsp transcriptional pathway (spa, SAR0143, clfA, SAR1014, SAR1745, ureA, ureG, SAR2427, fnbA, clfB, sasA, SAR2763) | 16/147 | 0/109 | 0.0001 *** |
| SarA transcriptional pathway (SAR0109, spa, SAR0211, pyrAA, SAR1397, agrC, agrA, SAR2245, SAR2420, SAR2430, hlgB, fnbA, arcC, sasA, lip) | 20/147 | 1/109 (agrC) | 0.0001 *** |
| AgrA transcriptional pathway (spa, SAR0211, pyrAA, SAR1397, sucA, SAR1466, hemL, agrC, agrA, SAR2430, hlgB, hlgC, clfB, arcC, sasA, lip) | 21/147 | 1/109 (agrC) | <0.0001 *** |
| Cell wall (spa, clfA, fnbA, clfB, sasA) | 9/156 | 0/115 | 0.01 * |
| Cell adhesion (clfA, fnbA, clfB) | 6/156 | 0/115 | 0.04 * |
| Pathogenesis (spa, SAR0115, SAR280, SAR0464, SAR0739, saeR, clfA, ebh, rot, SAR2035, SAR2448, hlgA, hlgB, fnbA, clfB, sasA) | 21/156 | 2/115 (ebh) | 0.0006** |

